# Genome-wide mapping of 5ʹ-aldehyde terminus induced by reactive oxygen species

**DOI:** 10.1101/2025.02.19.636396

**Authors:** Junzhou Wu, Yang Jiang, Jingjing Sun, Peter C. Dedon, Shana J. Sturla

## Abstract

DNA single strand breaks (SSBs) are abundant lesions due to cell metabolism and reactive oxygen species (ROS) and can lead to replication folk collapse, double strand break formation, genome rearrangements, cell death and disease. Among numerous chemical forms of SSBs, 5ʹ-aldehyde terminus are the most abundant generated by hydroxyl radicals and pose significant challenge for cellular repair machinery due to the lack of specific end processing process, potentially leading to more severe biological consequences than other readily repairable SSBs. Herein we developed a new strategy to locate 5ʹ-aldehyde terminus in genomic DNA at single-nucleotide resolution. The principle involves labelling the 5ʹ-aldehyde terminus with an aminooxy-functionalized oligonucleotide, giving rise to a biocompatible altered DNA linkage and allowing labelled sites to be amplified by polymerase chain reaction. We sequenced the 5ʹ-aldehyde terminus distribution in genomic DNA, nuclei, and cells following activation of the Fenton reaction. The results revealed a significant preference for adenine bases in DNA lesions and provided insights into the genome-wide distribution of such DNA damage, correlating with genomic features and chromatin accessibility. This method provide a new strategy for studies aiming to understand the biological and toxicological impacts of 5ʹ-aldehyde termini in DNA as the form species of single strand break induced by reactive oxygen species from a human genome.

## Introduction

Reactive oxygen species (ROS), including superoxide (O_2_^-^), hydrogen peroxide (H_2_O_2_), hydroxyl radicals (OH·), and singlet oxygen (^1^O_2_), are continuously produced in cells due to metabolism and external chemical exposures. Given their capacity to damage DNA, ROSs are major contributors to genetic instability and are implicated in various oxidative stress-induced diseases^1^. ROS can oxidise DNA bases forming, for example as 8-oxoguanine (8-oxoG), which is invoked in mutagenesis and carcinogenesis^2^. Additionally, 2-deoxyribose oxidation jeopardizes genetic stability and cell survival by inducing single- and double-strand DNA break (SSB and DSB)^3,4^. Exposure of the hydrogen atoms at the C5 position to diffusible molecules^5^ makes them particularly susceptible to abstraction by hydroxyl radicals, resulting in formation of a 5ʹ-aldehyde terminus (5ʹ-AT) and simultaneous DNA strand break^6-8^. Consequently, 5ʹ-AT represents one of the most prevalent oxidized 2-deoxyribose lesions^6^, possessing a half-life of over one week in free DNA under physical condition^8^. Because there is a lack of repair proteins for 5ʹ-AT end processing, it is believed to be repaired by the long patch base excision repair (BER) pathway with less efficiently compared to other DNA lesions that repaired by short patch BER pathway. The combined insights suggest that 5ʹ-AT may present significant challenges within cellular genome. However, the distribution and the biological implications of unrepaired 5ʹ-AT in a genome remain poorly understood, possibly due to a lack of high-sensitivity detection methods.

Mapping 5ʹ-AT at a genome scale is imperative for elucidating its formation and repair as well as its role in disease progression, yet current methodologies lack the precision required for accurate sequencing 5ʹ-AT. Traditional methods for analyzing aldehydic DNA lesions often utilize biotinylated reagents containing hydroxylamine groups, known as Aldehyde Reactive Probes (ARP), which has been applied for global quantification^9^ or for genome-wide mapping^10^ of abasic sites. However, the chemical probe lack the specificity required for 5’-AT detection, as it reacts with a wide range of aldehydic DNA lesions and modifications. Several next-generation sequencing (NGS) methods, such as S1 END-seq^11^, SSiNGLe-ILM^12^, GLOE-Seq^13^, and SSBlazer^14^, have focused primarily on enzymatic ligatable SSBs arising from mixed sources like oxidative lesions, DNA replication, and damage repair. Therefore, there remains a critical need for the development of methodology that can selectively detect and map ROS-induced SSBs across the genome to further understand its biological significance.

In this study, we present a chemical-specific approach to sequence 5ʹ-AT caused by ROS. 5ʹ-AT was labelled with a 3ʹ-aminooxy 5ʹ-biotin-modified oligodeoxynucleotide (ODN), resulting in a biocompatible oxime-linked DNA that can be read through by DNA polymerases and amplified by polymerase chain reaction (PCR). In this way, the ligation products generated from 5ʹ-AT can be sequenced by NGS, and the code sequence can be used as a readable marker to identify specific genome locations of 5ʹ-AT and create single-nucleotide resolution genomic maps. Herein, we generated a nucleotide-resolution map of 5ʹ-AT for the genome of a human foreskin fibroblast cell line. We evaluated genomic factors like sequence contents and genome accessibility to understand factors driving the distribution of 5ʹ-AT in the human genome.

## Results

### Labelling of aldehyde-modified oligonucleotides with the code sequence

To optimize labeling efficiency, we investigated the labeling reactions between 5ʹ-AT, abasic site modified ODNs and 3ʹ-aminooxy modified code sequence. Notably, AP sites exhibit another major aldehyde source in the genome and may also react with aminooxy group. Storing oligodeoxynucleotides (ODNs) containing aldehydic lesions (5ʹ-AT, abasic sites) and 3ʹ-aminooxy modifications is challenge due to their high reactivity with amines or aldehydes in solution, as well as their susceptible to cleavage. Therefore, all modified ODNs were stored as precursors and freshly prepared before use. This preparation included 5ʹ-AT containing ODNs generated *via* NaIO_4_ treatment of vicinal diol modified ODNs (**Diol-F, GFP-Diol**)^15^, 3ʹ-aminooxy-modified ODNs prepared by removing dimethoxytrityl protection at acidic condition (**Bar-ON-Btn**) (Table S2), and abasic site containing ODN produced by treating an uracil containing ODN (**T21F-U**) with uracil-DNA glycosylase (UDG) (Figure S2). The labeling reaction for both 5ʹ-AT and abasic site was conducted under mildly acidic condition (pH 5.4) with an excessive amount of 3ʹ-aminooxy code sequence to facilitate the rapid oxime conjugation. Within 2 hours, the 5ʹ-AT labeling reaction was fully completed, yielding a 35-mer conjugated product, as evidenced in the denaturing polyacrylamide gel (Figure 1b). In contrast, the labelling reaction between abasic site and 3ʹ-aminooxy code sequence only yielded 7% conjugated product and formed 74% backbone breakage products through β-elimination at the abasic site (Figure S2). These results demonstrate that 5ʹ-AT in a DNA sequence can be efficiently labelled by a 3ʹ-aminooxy code sequence under a mild acidic condition, although the abasic site undergoes labeling with significantly reduced efficiency.

**Figure 1.**
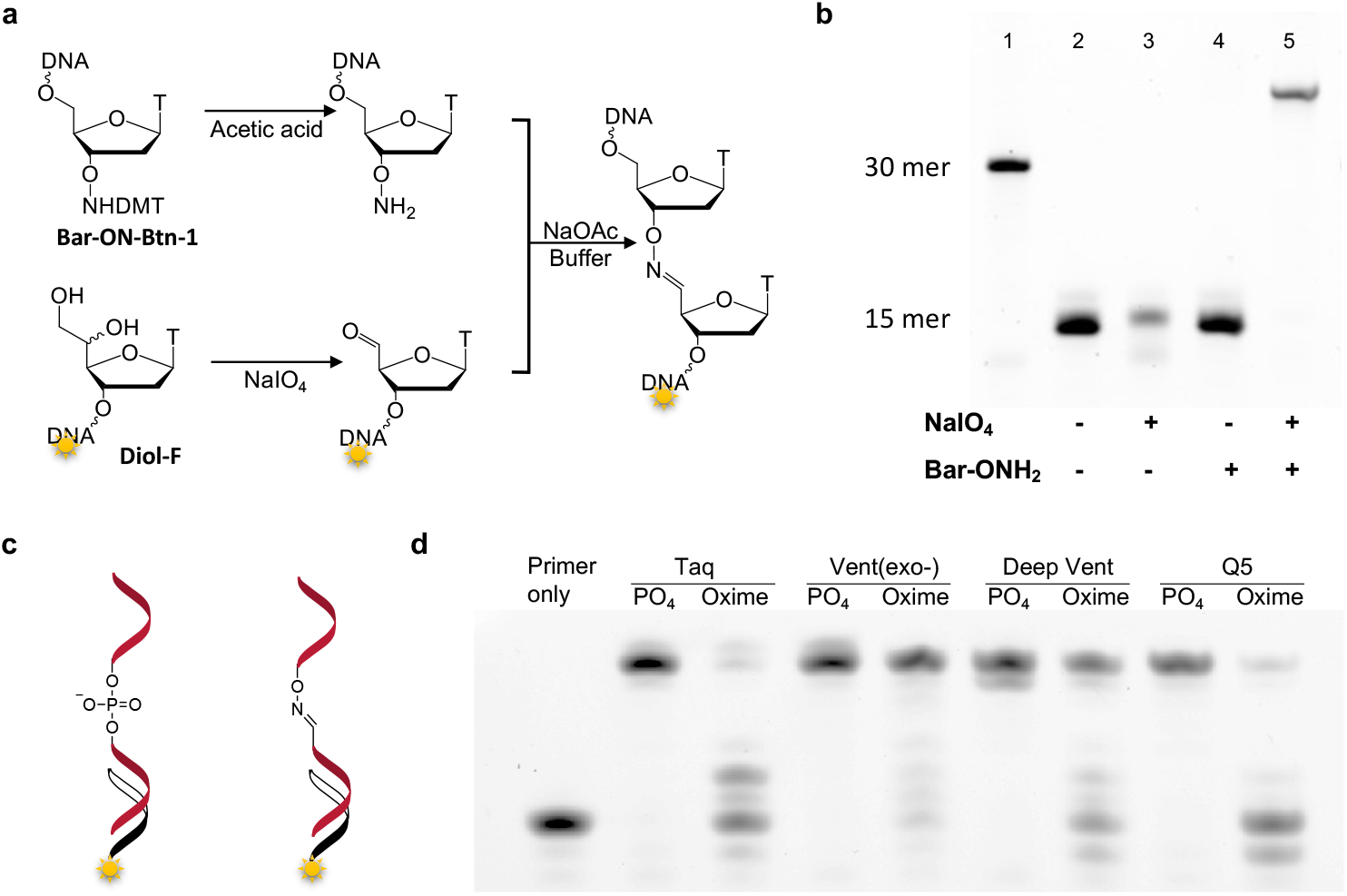
(a) Labelling of aldehyde modified oligonucleotide (Diol-F) with code sequence (Bar-ON-Btn-1). (b) Denaturing PAGE analysis of products after labelling reactions. Only fluorophore labelled 5ʹ-AT containing ODN is visible. Lane 1 shows a 30-mer ODN marker. The slower migrating product in Lane 5 is attributed to conjugated oxime linkage ODN. (c) Bypass of a DNA template containing a site-specific oxime linkage (lanes 3, 5, 7, 9, 11) or identical template without modification (lanes 2, 4, 6, 8, 10) by different DNA polymerases. Lane 1 indicates a control without adding polymerase.

### Bypass and amplification of ligated products raised from aldehyde labeling

A cornerstone of being able to sequence 5ʹ-AT is the choice of a DNA polymerase that can amplify the labelled product resulting from the reaction of 5ʹ-AT with 3ʹ-aminooxy modified code sequence, rather than from other aldehyde-modified nucleotides. We initially tested the polymerase bypass of an ODN (**T21-ON**) that contains a oxime linkage - the reaction product between 5ʹ-aldehyde and 3ʹ-aminooxy -instead of a canonical phosphodiester linkage(Figure 1c, S1). All four tested polymerases were able to bypass the oxime linkage, yielding full-extension products that were identical to those from canonical ODNs, without any insertions or deletions at the modification site (Figure 1d). Furthermore, Vent (exo-) and Deep vent bypassed the oxime linkage with more than 50% efficiency. These indicate that replacement of the phosphodiester bond in DNA by an oxime linkage is a viable DNA synthesis template.

The next step is to identify a polymerase that can selectively amplify the labeled product from 5ʹ-AT instead of AP sites. First, 0.9-kb dsDNAs containing either a vicinal diol or a deoxyuridine modification were prepared by ligating gapped dsDNA with synthetic complementary ODNs (**GFP-diol** or **GFP-U**) (Figure S3). Subsequently, the 5ʹ-AT or AP site was generated by treating the dsDNA with NaIO_4_ or UDG. They were labelled with 3ʹ-aminooxy code sequence, then both were submitted for PCR reactions with one primer for dsDNA and one primer specific for the code sequence. We observed that Taq, Deep vent, and Q5 polymerases could amplify the labeled 5ʹ-AT specifically (Figure 2a), while Vent (exo-) did not differentiate 5ʹ-AT and AP site, as it yielded amplicons from both. Sequencing the 5ʹ-AT amplicon revealed the correct bypass of the T-oxime-T structure, evidenced by AA dimer in amplicons (Figure 2b). Thus, Taq, Deep vent and Q5 amplified oxime-labeled 5ʹ-AT without interference from the AP site,.

**Figure 2.**
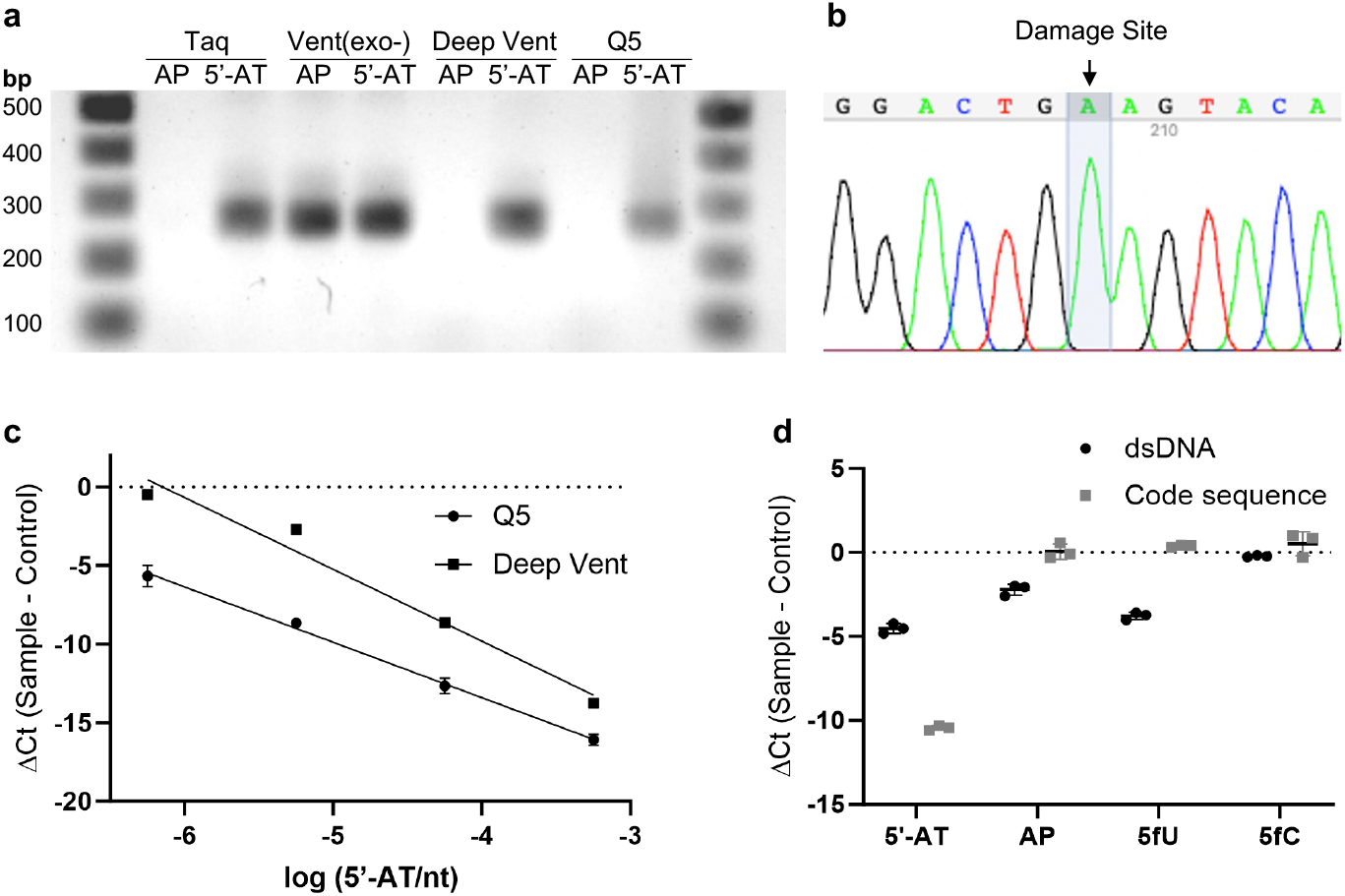
(a) Amplification of labelling products from abasic site (lanes 2, 4, 6, 8) or 5ʹ-AT (Lanes 3, 5, 7, 9) by different DNA polymerases. (b) Sanger sequencing data from 5ʹ-aldyhyde amplicon showing code sequence marked 5ʹ-AT lesion. (c) qPCR Ct values using Q5 or Deep vent polymerase as a function of relative 5ʹ-AT concentrations (number of 5ʹ-AT/10^6^ nt).

### 5ʹ-AT enrichment and detection sensitivity

To assess whether the labeling and bypass strategy was suitably sensitive not only for sensing 5ʹ-AT in ODNs, but also at ultra-low levels anticipated in genomic DNA, we mixed varying amounts of 5ʹ-AT modified dsDNA with native dsDNA, producing reference dsDNA mixtures with 5ʹ-AT quantities ranging from 0.56 to 560 lesions per 10^6^ nt (Table S4). Following labeling with 3ʹ-aminooxy code sequence, all samples were quantified by qPCR with high-fidelity DNA polymerase, either Deep vent or Q5. At the lowest test 5ʹ-AT input (0.56/10^6^ nt), the ΔCt value relative to canonical dsDNA for Q5 is -5.7 and only -0.5 for Deep vent (Figure 2c). The ten-fold serial dilutions of 5ʹ-AT demonstrated a linear response over four logs for Q5 (R^2^ = 0.99, Efficiency = 92%) and three logs for Deep vent (R^2^ = 1.00, Efficiency = 52%, excluding the data with the lowest 5ʹ-AT). Thus Q5 polymerase is both efficient and sensitive for detecting 5ʹ-AT concentrations as low as 0.56 lesions per 10^6^ nt, consistent with cell-relevant damage levels.

During NGS library preparation, a 5ʹ biotin-modified code sequence was employed to enrich labeled fragments, thereby further reducing background noise. This approach was first tested using dsDNA fragments contain four types of common aldehyde modificatiosn in DNA, i.e. 5ʹ-AT, AP site, 5-formyl-deoxyuridine (5fU) and 5-formyl-deoxycytidine (5fC). After labelling reaction with code sequence, the dsDNAs were enriched on streptavidin beads and subjected for qPCR with one set of primers targeted dsDNA and another set of primers targeted code sequence (Figure S5). The amplicon of dsDNA was detected in 5ʹ-AT, AP and 5fU samples, while the amplicon of code sequence only observed in 5ʹ-AT sample (Figure 2d). Those findings suggest that dsDNAs containing 5ʹ-AT, AP site and 5fU could be labeled by code sequence and enriched, while only the oxime linkage raised from 5ʹ-AT could be bypassed and amplified as we expected. The results also suggest that the aldehyde group in 5fC is not as reactive as 5ʹ-AT, AP site and 5fU^10^. In summary, after both enrichment and bypass, a highly selectivity amplification was achieved for the 5ʹ-AT lesions in comparison to other DNA aldehyde damage/modifications.

### A genome-wide map of 5ʹ-AT

Having established an efficient and sensitive method to label and amplify 5ʹ-AT, we used this novel approach termed “5AT-seq” for genome-wide sequencing of 5ʹ-AT with single-nucleotide resolution (Figure 3a). Thus, we applied it to BJ-5ta cells, an immortalized primary fibroblast cell line, to investigate DNA damage and repair in a non-cancerous system. First, BJ-5ta genomic DNA containing 5ʹ-AT was labeled with 3ʹ-aminooxy code sequences (**Bar-ON-mix**), and then sheared by sonication into short fragments to preempt sonication-induced 5ʹ-AT strand breaks. The labeled DNA was then denatured and enriched using streptavidin beads. Then, the enriched DNA was eluted from beads based on previous protocol^16^ and ligated with P7 adapter (Table S2). The ligation product was subjected to a 5-cycle bypass PCR using Q5 polymerase, followed by an indexing PCR of the purified bypass product, and then submitted for Illumina sequencing (Figure 3a). In the sequencing data analysis, the first 5 bases of read 2, derived from the Bar-ON-mix code sequences, were used to filter out unspecific read, and then trimmed prior to alignment and further data analysis.

**Figure 3.**
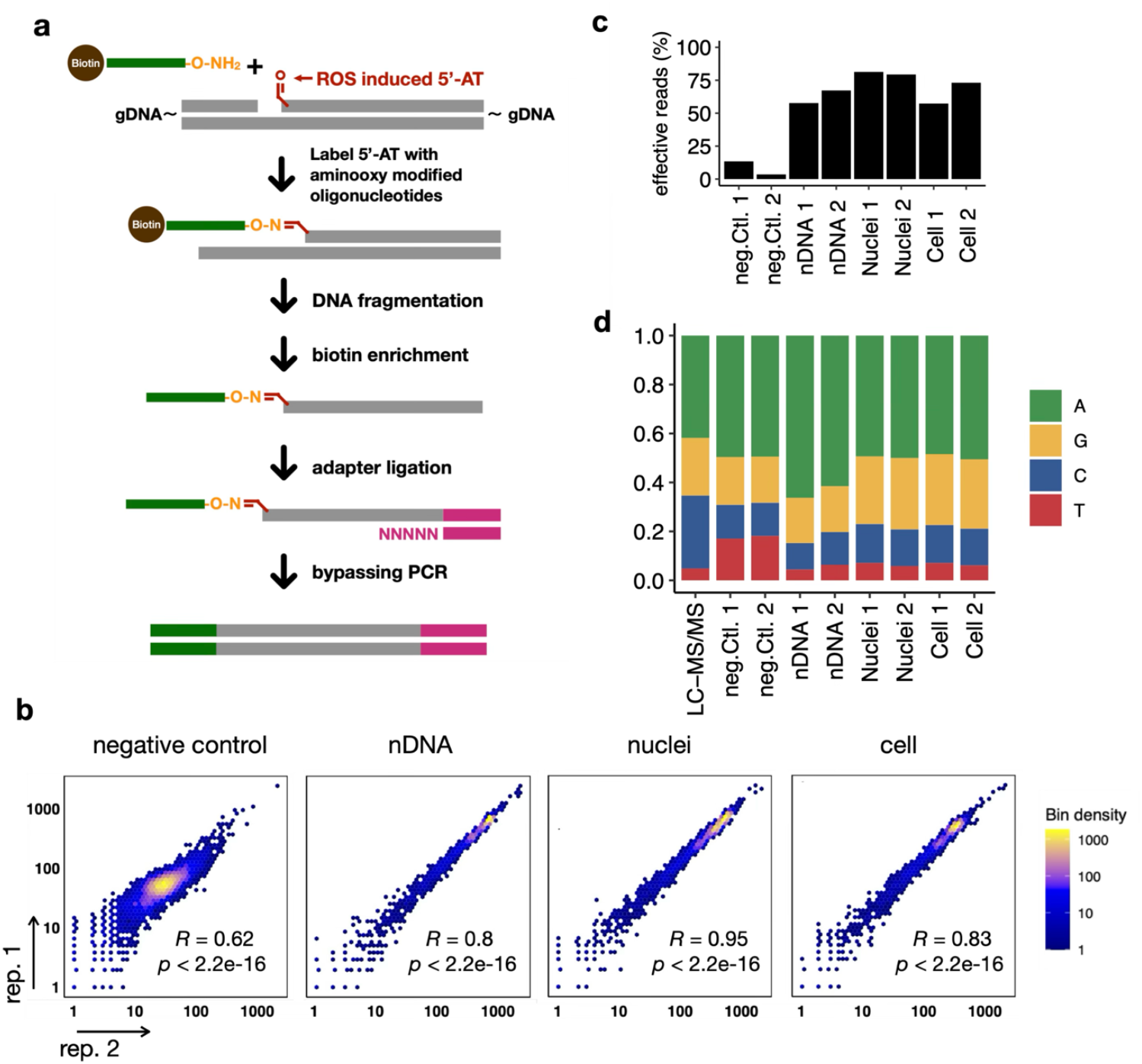
Schematic diagram of 5AT-seq and preliminary data analysis. (a) Schematic representation of the method. (b) Scatter plots showing the Spearman’s correlation of biological duplicates at 100 kb bins. (c) Unique, high-quality reads in percentage of total sequencing depth. (d) Base composition of 5ʹ-AT sites quantified by LC-MS/MS (lane 1) and sequencing (lane 2-9), showing a preferred 5ʹ-AT formation on DNA bases.

### High reproducibility and specificity of 5AT-seq Results

To test how nucleosome packaging^17-20^ may act as a barrier of DNA for ROS-induced 5ʹ-AT formation, we exposed naked DNA, nuclei, and cells to extracellular ROS generated via the Fenton reaction for 5AT-seq. In treated cells, we identified a higher ratio of effective (unique, high-quality) reads at 65.2% in average, compared to 8.5% in untreated cells (negative control) (Figure 3c). This result aligns with qPCR analysis, which indicates approximately an 10-fold increase (ΔCt = -4.0) in 5ʹ-AT lesions in treated cells relative to untreated cells. This observation underscores the low abundance of 5ʹ-TA in the negative control and the high specificity of 5AT-seq. Then, we assessed the correlation among sequencing libraries replicate (n = 2), and observed high Spearman’s correlation coefficients between duplicate samples across 5 kb – 100 kb bins, confirming the reproducibility of 5AT-seq (Figure 3b, S6a). Both intragroup and intergroup comparisons showed higher correlations for naked DNA, nuclei, and cells compared to the negative control (Figure S6a, S6b). Additionally, by principal component analysis, there was clear clustering of duplicates, and data from each experimental condition separated along the first component, pointing to distinctive genomic distributions of 5ʹ-AT under varying conditions (Figure S6c). These findings demonstrated a strong similarity in 5ʹ-AT patterns among treated samples, which were distinctly different from the non-treated negative control, highlighting the reproducibility and specificity of 5AT-seq.

### Base preferences of 5ʹ-AT

To investigate the sequence preference of ROS-induced lesions, we analyzed the base composition at 5ʹ-AT sites from all samples. We found a preferred base composition at the strand break locations (Figure 3d). In the treated nDNA, the ratio of 5ʹ-AT production on individual bases is 12:3:2:1 (A:G:C:T), with a pronounced preference for A. This A preference was further validated using LC-MS/MS analysis. Comparing to nDNA, treated cells and nuclei exhibited a noticeable increase of lesions on G and C, which was later identified to be due to a preferred damage accumulation in GC rich regions. Notably, the base composition in unexposed cells was similar to exposed cells and nuclei, but with a slight elevation on T, which possibly represents the steady state 5ʹ-AT lesions under an equilibrium between formation and repair in BJ-5ta cells.

### Genome-wide distribution of 5ʹ-AT

Having examined the lesion pattern at single-base resolution, we then explored the genomic distribution of 5ʹ-AT. By using chr8 as an example, we found 5ʹ-AT distribution through the chromosome (100-kb bins) varied among samples (Figure 4a), such that for exposed nDNA, 5ʹ-AT was evenly distributed throughout the chromosome, whereas for both nuclei DNA and cellular DNA, whether exposed or unexposed, 5ʹ-AT levels fluctuated consistent with an influence of chromatin structures. To delve deeper into the potential influence of specific genomic features, we compared 5ʹ-AT profiles with GC content, chromatin accessibility, euchromatin and heterochromatin markers, and transcription factor binding sites^21-24^. The spearman correlation coefficient was calculated between the 5ʹ-AT count and the genomic feature coverage in 100-kb bins. We observed that the distribution of 5ʹ-AT in nDNA was predominantly negatively correlated with most of the examined features, most notably with GC content (R=-0.48, Figure 4b). Given that 5ʹ-AT is prone to form on A, it is not surprising to see the negative correlation between 5ʹ-AT in nDNA and GC content, along with other features exhibiting GC preference like H3K4me3^25^ and CTCF^26^. In contrast, the distribution of 5ʹ-AT in untreated cells, and treated nuclei and cells was generally positively correlated with those genomic features, notably with DHS, H3K27ac and the FOXA binding sites (Figure 4b), which are typically found in active genomic regions. DNase-seq and ATAC-seq data directly represent chromatin accessibility; H3K27ac and H3K4me3 are highly enriched in the promoter region of actively transcribed genes^27^; and GATA4 and FOXA binding sites are typically found in accessible genomic regions^24^. This high accessibility makes these regions more susceptible to DNA damage during exposure^28^, explaining their positive correlation with 5ʹ-AT distribution. Furthermore, the positive correlation between GC content and active genomic features (Figure S7) explains the elevated damage ratio on G and C in the treated nuclei and cells (Figure 3c). To provide a comprehensive overview, counts of 5ʹ-AT and DHS in 100-kb bins were plotted across the individual chromosomes of the BJ5 cell line (Figure S8), which clearly highlights the relationship between 5ʹ-AT enrichment and genome accessibility. Notably, for two heterochromatin indicators that linked with low genome accessibility^24,28,29^, H3K9me3 and H3K27me3, there were no significant association with 5ʹ-AT (Figure 4b). We speculate that the weak correlation is due to the low-sampling of the histone mark ChIP-Seq data (Table S1), which is not sufficient for a genome-wide scale analysis.

**Figure 4.**
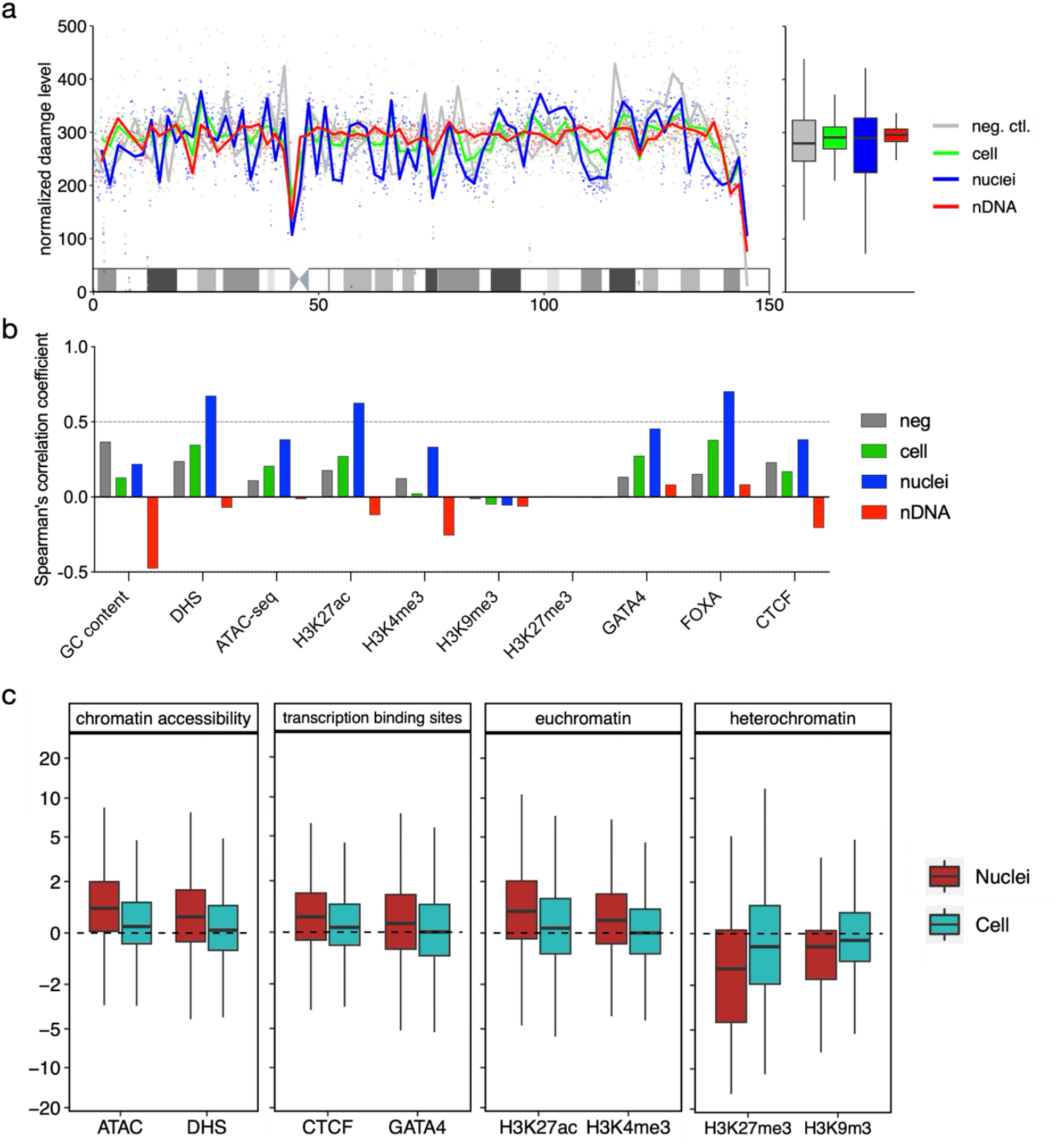
(a) Detailed view of 5ʹ-AT distribution on Chromosome 8. Plot lines represent the average 5ʹ-AT level calculated from two independent experiments. The LOESS curve is fitted to the damage level based on 100-kb bins along Chromosome 8. The boxplot on the right side is plotted using the same data from the chromosome view. (b) Spearman’s correlation coefficients for the correlation of 5ʹ-AT levels with genomic features (100 kb resolution). Data represent the mean calculated from two independent experiments. (c) Boxplots represent the relative 5ʹ-AT levels in the selected genomic features. Source data for genomic feature annotation are listed in Table S2.

To further investigate the factors influencing 5’-AT distribution, we assessed its relative abundance across genomic features in treated nuclei and cell samples, using treated nDNA as a baseline. Negative controls were excluded due to insufficient reads (n < 1 million). A positive differential indicated 5ʹ-AT enrichment, while a negative value signified depletion. The analysis revealed significant 5ʹ-AT enrichment in accessible regions, including euchromatin and transcription factor binding sites (e.g., GATA4, FOXA1), and depletion in heterochromatic regions marked by H3K27me3 and H3K9me3 (Figure 4c). This suggests 5ʹ-AT formation occurs primarily in euchromatin, with low protein interference and increased ROS susceptibility.

## Discussion

Here, we introduced 5AT-seq, a novel method for mapping 5’-AT, a prevalent form of DNA damage induced by ROS, across the human genome. The 5AT-seq strategy involves the labelling of 5’-AT sites with a code sequence using an aminooxy-functionalised oligonucleotide, thereby enabling site-specific amplification by PCR and subsequent sequencing at single-nucleotide resolution. Notably, 5AT-seq is capable of distinguishing 5’-AT lesions from abasic sites, which a major source of aldehydes in the genome. We validated the method was by testing it on naked DNA, nuclei, and whole cells subjected to the Fenton reaction, demonstrating its high specificity and reproducibility. Furthermore, a genome-wide analysis of 5’-AT distribution was performed, uncovering its associations with key genomic features.

Our analysis at the nucleotide level revealed a marked preference for formation of 5-AT at positions of deoxyadenosine, accounting for approximately 50% to 60% of C5ʹ oxidation events. This observation was corroborated by LC-MS/MS analysis of methoxylamine-derivatized 5ʹ-AT lesions. However, it’s unlikely that hydroxyl radical could distinguish between different C5ʹ of dA, dG, dC, dT. Considering the brief lifetime and limited travel distance of the hydroxyl radicals^30^, we hypothesize that the ferric·EDTA complex may have sequence-binding preference or altered catalytic activity upon interaction with DNA sequences, leading to a higher concentration of the racicals at specific sequence, i.e. A rich sequence in our studies. Previous study has pointed out that ferric ion could form strong stacking interactions with purine nitrogen atoms^31^, potentially explaining in part the preferential formation of 5ʹ-AT on dA/dG over dC/dT. In another study, linear dichroism signals during ferric·EDTA induced DNA cleavage indicated the resistance of poly(dG-dC) against Fenton reaction compared to poly(dA-dT) and poly(I-dC). The author suggested that ferric·EDTA may bind at the N^2^ amino group of G in the minor groove and thereby suppressing the action of the reactive oxygen species on DNA backbones^32^. Furthermore, GC rich sequence may form sinks for hydroxyl radicals due to the C8 of guanosine^33,34^, producing the most abundant oxidative base lesion 8-oxoG instead of 5ʹ-AT. Therefore, we suggest that ferric-· EDTA has a preferential binding affinity for the N^7^ position of dA, facilitating ROS reacting with the C5ʹ position of dA. Conversely, when ferric-EDTA is bound to the N^2^ position of dG, ROSs are less likely to react with the C5ʹ due to the increased travel distance of ROSs from N^2^ to C5ʹ. Alternatively, binding of ferric-EDTA to the N^7^ position of dG permits ROS to react with both the C8 and C5ʹ positions, thereby reducing the formation of 5ʹ-AT. Further studies are required to provide more insights into the mechanisms of dA preference of 5ʹ-AT formation by Fenton reaction.

It is worth noting that higher correlation coefficients between 5ʹ-AT and genomic accessibility were observed in the treated nuclei compared to the treated cells. We speculate that this is due to the absence of cellular activity after nucleus isolation, resulting in a relatively stable chromatin structure.^35^ In live cells, on the other hand, chromatin structure is dynamic,^36,37^ especially when the cell cycle is not synchronized, which is the case in this study. Therefore, chromatin within these cells is expected to be more accessible for the chemical exposure, leading to the DNA damage to be more evenly distributed along the genome. Additionally, the repair of SSB is generally rapid with a half-time of a few minutes in normal cells^38,39^, thus, a certain level of 5ʹ-AT would be repaired before DNA extraction. Considering the relatively high repair capacity in the open genomic regions^20,40^, the variation of damage levels related with chromatin structure, could be mitigated in cells.

In conclusion, we created 5AT-seq, a novel approach for single nucleotide resolution sequencing of 5ʹ-AT lesions induced by ROS in human genome DNA. Given that 5ʹ-AT is a primary outcome of 2-deoxyribose oxidation, our sequencing results may serve as a proxy for single-strand breaks attributable to ROS activity. The results have demonstrated a distinct preference for adenine and a strong correlation with chromatin accessibility of 5ʹ-AT. Future research will focus on exploring the dynamics of damage formation and repair over time or in repair-deficient cells, conducting mechanistic studies on adenine preference, and assessing the impact of hydroxyl radical sources—such as γ-radiation and enediyne antitumor antibiotics—on the distribution of 5ʹ-AT. With these studies, the 5AT-seq method will enhance our understanding of their biological effects and the broader implications of oxidative DNA damage on cellular processes.

## Materials and Methods

### Materials

Oligonucleotides and indexing primers were purchased from Eurogentec. BJ-5ta human skin fibroblast cells were from ATCC (CRL-4001) and regularly tested. Genomic DNA was extracted using the Monarch Genomic DNA Purification Kit (New England Biolabs, NEB) according to the manufacturer’s instructions. DNA concentrations were measured using a Quantus fluorometer with QuantiFluor ONE dsDNA Dye (Promega). All chemicals for solid phase synthesis of oligonucleotides were purchased from Glen Research. All other chemicals were purchased from Sigma Aldrich.

### Modified oligonucleotide synthesis

Diol-F, GFP-diol, Bar-ON-Btn and T21-ON modified ODNs were synthesized on 1 μmol scale on a MerMade 4 Oligonucleotide synthesizer (BioAutomation Corporation). For the coupling step, 5-Ethylthio-1H-tetrazole (ETT) was used as activator (0.5 M in anhydrous CH_3_CN), with a coupling time of 2 × 15 s for standard nucleoside phosphoramidites (0.1 M in anhydrous CH_3_CN) and 4 × 30 s for phosphoramidites **9** and **14** (0.1 M in anhydrous CH_3_CN). The capping step was performed with acetic anhydride by using commercial solutions (Cap A: Ac_2_O/pyridine/THF, 10:10:80, v/v/v; Cap B: 10 % N-methylimidazole in THF) for 30 s. Oxidation was performed for 15 s with 0.1 M Iodine in THF/pyridine/water (78:20:2). Detritylation was performed with 3 % trichloroacetic acid (TCA) in Dichloromethane (DCM) for 2 × 30 s. Final trityl groups were remained (DMT-on) for all ODNs. For Dio-F, standard DNA phosphoramidites (Glen Research) and phosphoramidite **9** were used. For Bar-ON-Btn ODNs, 5’ -> 3’ synthesis phosphoramidites (Glen Research) and phosphoramidite **14** were used. For T21-ON, the sequence was first synthesized until the modification site using standard DNA phosphoramidites (Figure S1). After detritylation, CPG-linked ODN were stirred in a 0.5 M N, N’-dicyclohexylcarbodiimide in DMSO (with 1.7% dichloroacetic acid) mixture for 30 min at ambient temperature to form 5ʹ-AT. The solvent was removed and the CPG was first washed twice with DMSO and twice with acetonitrile. Then 3ʹ-aminooxy modified thymidine (compound **12**, 20mg), 250 μL methanol (with 0.1% acetic acid) were added to the CPG. The mixture was rotated overnight at ambient temperature. The solvent was then removed and the CPG was washed twicewith methanol and twice with DCM. Then, 5ʹ-*tert*-butyldimethylsilyl (TBDMS) protecting group on the modified nucleotide was removed by treating the CPG with tetrabutylammonium fluoride hydrate (1M in THF) for 4 hours. The solvent was removed and the CPG was washed twice with dry THF and twice with dry acetonitrile. Then, the sequence after oxime modification was synthesized automatically using standard DNA phosphoramidites on a Mermade 4 Oligonucleotide synthesizer.

The deprotection of modified ODNs was carried out with 30% aqueous NH_4_OH at ambient temperature for 16 hours. After decanting the supernatant from the CPG support, 50 μL of triethylamine (TEA) was added. The mixture was concentrated to dryness in a MiVac centrifugal evaporator (GeneVac). The ODNs were purified by HPLC (Agilent 1200 Series) using a Phenomenex Luna C18 250 × 4.6 mm column with a gradient of acetonitrile (10% to 40% buffer B over 20 min, flow rate 1 mL/min), buffer A: 0.05 M triethylammonium bicarbonate buffer, pH 7.5, buffer B: acetonitrile. Elution was monitored by UV absorption at 260 nm. After HPLC purification, ODNs were concentrated to dryness in a MiVac centrifugal evaporator. Bar-ON-Btn ODNs were resuspended in deionized water for further use. The 5ʹ-DMT group on purified ODNs were removed by 80% acetic acid for 30 min at room temperature. Acetic acid was removed with a MiVac centrifugal evaporator. The ODNs were desalted by Sep-Pak classic C18 cartridges, concentrated and re-dissolved in deionized water. The composition was confirmed by direct-injection mass spectrometric analysis with an Thermo LTQ Velos.

### Labelling of a 5ʹ-aldehyde-modified oligonucleotide with a code sequence

A 5ʹ-aldehyde modified ODN was prepared by reacting a diol-modified ODN (Diol-F, 2 µM) with 25 mM NaIO_4_ in 100 mM NaOAc (20 µL, pH 6.0) at room temperature for 60 min. The ODN was desalted by passing through a micro bio-spin 6 column (Bio-Rad). An aminooxy-modified ODN without DMT protection was prepared freshly every time before use by treating Bar-ON-Btn (1 mM, 1 µL) with acetic acid (9 µL) at room temperature for 30 min. Acetic acid was removed by a MiVac centrifugal evaporator. Then, 18 µL 5ʹ-aldehyde modified ODN and 2 µL NaOAc (1 M, pH 6.0) were added into dried aminooxy modified ODN. The reaction mixture was incubated at room temperature for 2 hours. The reaction was quenched with formamide loading buffer and analyzed by denaturing urea polyacrylamide gel electrophoresis (urea-PAGE). Gels were imaged with a ChemiDoc XRS+ System (Bio-Rad). Band intensities were quantified with Image Lab (Bio-Rad).

### Labelling of an oligonucleotide containing abasic site with the code sequence

An ODN containing abasic site was prepared with 3 µM T21F-U, 2 μL UDG (NEB, M0280S) in 100 μL 1 × UDG buffer at 37 °C for 1 h. The resulting ODN was purified with Monarch PCR & DNA Cleanup Kit (NEB) using a protocol for ODNs. The labelling reaction was carried out between the ODN containing abasic site and the aminooxy modified ODNs as described above for the 5ʹ-aldehyde modified ODN. The reaction was quenched with formamide loading buffer and analyzed by urea-PAGE. Gels were imaged with a ChemiDoc XRS+ System. Band intensities were quantified with Image Lab.

### Bypass of oxime-linked ODNs

The bypass reaction (20 μL) contained 2 µM oxime linker containing ODN as template (T21-ON), 1 µM P14F as primer, 250 μM dNTP mixture and 1x supplied reaction buffer for each polymerase, i.e. standard taq buffer for Taq polymerase (NEB, M0273S), ThermoPol buffer for Vent(exo-) (NEB, M0257S) and Deep vent (NEB, M0258S), and Q5 reaction buffer for Q5 polymerase (NEB, M0491S). The reaction mixtures were heated up to 80 °C and cooled down to 50 °C at 1 °C/minute speed. Then, 1 unit of each polymerase was added to the reaction mixture and keep at 50 °C for 30 minutes. The reaction was quenched with formamide loading buffer and analyzed by denaturing urea polyacrylamide gel electrophoresis (urea-PAGE).

### Construction and amplification of dsDNA containing modifications

dsDNA fragments containing a site-specific modification was first prepared by ligation of a short modified ODN and a 0.9-kb-gapped DNA duplex. In brief, a pEGFP-W1 plasmid was constructed to contain two Nb.BbvCI (NEB, R0631S), a nicking endonuclease, cleavage sites. A 0.9 kb DNA duplex was amplified from the plasmid with primers GFP-Pr409 and GFP-Pr1296 and Taq DNA polymerase, and then subjected to Nb.BbvCI digestion. A complimentary ODN was added to aid the removal of cutted short sequence and the gapped dsDNA was purified with Monarch Nucleic Acid Purification Kit. Then, the gap was filled by annealing of the dsDNA with GFP-Diol or GFP-AP in large excess. Nick sites were ligated by T4 DNA ligase (NEB, M0202S) at 16 °C for 4 h and the ligation product was purified with Monarch Nucleic Acid Purification Kit.

To perform PCR amplification of modified dsDNA, various amounts of dsDNA containing 5ʹ-AT were mixed with native dsDNA. The final 5ʹ-AT amount is ranged from 0.56-560 lesion/10^6^ nt (Figure S4). After the labelling reaction with Bar-ON-mix, excess ODNs was removed using Monarch Nucleic Acid Purification Kit, then the labelled dsDNA was pulled down using streptavidin beads following the manufacturersʹ instruction. The pulled down product was then subjected for PCR amplification. The primers used for amplification are Ex-Bar-mix and GFP-Pr409, Ex-Bar-mix was prepared by mixing equal amount of Ex-Bar-1 to 4 to the desired concentration. The PCR was performed under the following condition: 60 s at 98 °C, (10 s at 98 °C, 180 s at 65 °C) x3, (10 s at 98 °C, 75 s at 65 °C) x35, hold at 4 °C.

### BJ-5ta cell culture and chemical exposure

BJ-5ta cells were cultured in medium containing a 4:1 mixture of DMEM (Thermo Fisher Scientific, 11965092): M199 (Thermo Fisher Scientific, 11150059) and 20% FBS with 10 mg/L Hygromycin B at 37 °C in a humidified atmosphere with 5% CO2. BJ-5ta cells grown in 10 cm dishes were washed with PBS then treated with 10 mL freshly prepared exposure PBS solution containing 2 mM Fe^2+^-EDTA, 10 mM H_2_O_2_, and 20 mM sodium ascorbate for 30 min under the incubation condition. After exposure, cell pellets were harvested by scrapping and immediately used for DNA extraction and eluted using 35 μL Milli-Q water. The eluted DNA was immediately used for library preparation.

### BJ-5ta cell nuclei isolation and chemical exposure

Nuclei were isolated as previously described.^42^ Briefly, BJ-5ta cells grown in 10 cm dishes were washed with ice-cold PBS and harvested by scrapping using 1 mL PBS. The scrapped cells were transferred to a 1.5 mL tube and centrifuge at 10,000 x g for 5 s. The supernatant was removed, and the cell pellet was resuspended in 900 μL ice-cold PBS containing 0.1% NP-40 (NP40S, Merck KGaA, Germany) by pipetting, then the tube was centrifuged at 10,000 x g for 5 s. Repeated the resuspension and centrifuge steps described above, and the nuclei pellet was resuspended in 150 μL PBS. The exposure was performed by mixing isolated nuclei (70 μL) with 10 μL sodium ascorbate (10 mM), 10 μL Fe^2+^-EDTA (1 mM), and 10 μL H_2_O_2_ (10 mM) in a 1.5 mL tube. The mixture then was incubated at for 30 min at 24 °C on a tube revolver (Thermo Fisher Scientific, 88881001) with oscillation mode. DNA was immediately extracted from the exposed nuclei and eluted from the silica column using 35 μL Milli-Q water. The eluted DNA was immediately used for library preparation.

### Chemical exposure of naked DNA

Extracted DNA from unexposed BJ-5ta cells (5 μg, 70 μL) was mixed with 10 μL sodium ascorbate (10 mM), 10 μL Fe^2+^-EDTA (1 mM), and 10 μL H_2_O_2_ (10 mM) in a 1.5 mL tube. The mixture was then incubated for 30 min at 24 °C on a tube revolver (Thermo Fisher Scientific, 88881001) with oscillation mode. The exposed naked DNA was purified by ethanol precipitation using 10 μL sodium acetate (3 M) and 250 μL ethanol. Purified DNA was resuspended using 18 μL Milli-Q water, and immediately used for library preparation.

### 5ʹ-AT labeling and enzymatic hydrolysis of DNA for LC-MS/MS

Genomic DNA (5 μg) in 70 μL PBS was mixed with 10 μL sodium acetate (1M, pH 5.4) and 20 μL methoxyamine (1M, pH 5.4). The mixture was incubated at 24 °C on a tube revolver with oscillation mode for 2 hours. The mixture was then purified by ethanol precipitation using 250 μL ethanol. The purified DNA were resuspended and hydrolyzed in a 50 µL digestion cocktail containing 5 U benzonase, 2 U calf intestinal alkaline phosphatase, 0.5 U phosphodiesterase I, 0.1 mM deferoxamine, 0.1 mM butylated hydroxytoluene, 5 ng coformycin, 2 mM MgCl_2_ and 10 mM Tris buffer pH 8.0. The mixture was incubated at 37 °C for 6 hours. Digestion enzymes were subsequently removed by passing mixture through a 10k Da MWCO spin filter. DNA samples were analyzed on a Phenomenex Synergi Fusion-RP C18 column (100 × 2.0 mm, 2.5 µm) coupled to an Agilent 1290 HPLC system and an Agilent 6495 triple quadrupole mass spectrometer. The LC system was conducted at 25 °C and a flow rate of 0.35 ml/min, with a gradient starting with 100% solution A (0.1% formic acid in H2O), followed by 0-3 min, 0%-5% solution B (0.1% formic acid in 70% acetonitrile); 3-13 min, 5%-25% solution B; 13-14 min, 25%-80% solution B; 14-16 min, 80% solution B. The LC-MS system used an electrospray ionization source in positive mode with the following parameters: gas temperature, 200 °C; gas flow, 11 L/min; nebulizer, 40 psi; sheath gas temperature, 250 °C; sheath gas flow, 12 L/min; capillary voltage, 1500 V. Multiple Reaction Monitoring (MRM) mode was used for detection of product ions derived from the precursor ions with the collision energy (CE) optimized for maximal sensitivity. Parameters for MRM detection: dA, 252/136/14 (precursor mass/product mass/CE(V)); dT, 243/127/4; dC, 228/112/8; dG, 268/152/8; MA-dA, 279/136/14; MA-dT, 270/127/4; MA-dC, 255/112/8; MA-dG, 295/152/8.

### 5AT-seq library preparation

Oligonucleotides used in library preparation are listed in Table S2. L-P7 adaptor (40 µM) were prepared by mixing equal volumes (20 µL) of 100 µM L-P7-5com and L-P7-5end with 10 µL 5x Annealing buffer (50 mM Tris, pH 8.0, 250 mM NaCl, 5 mM EDTA), heating to 98 °C, then slowly cool to 25 °C. Bar-ON-mix labelling oligo was prepared by mixing equal amount of 1 mM Bar-ON-Btn-1, Bar-ON-Btn-2, Bar-ON-Btn-3, and Bar-ON-Btn-4 oligos. Ex-Bar-mix primer (10 μM) was prepared by mixing equal amount of 10 μM Ex-Bar-1, Ex-Bar-2, Ex-Bar-3, and Ex-Bar-4 oligos.

Labelling oligo Bar-ON-mix (1 mM) was first deprotected by mixing 1 μL oligo with 9 μL acetic acid and incubate 20 min at 24 °C on a tube revolver with oscillation mode. The oligo was dried by centrifuge under vacuum for 5 min using a miVac Centrifugal Concentrator and resuspended using 2 μL sodium acetate solution (1M, pH=5.4) by thorough vortexing. Next, genomic DNA in Milli-Q water (18 μL) was added to the oligo, mixed by pipetting and incubated for 3 h at 24 °C on a tube revolver with oscillation mode. The labelled DNA was then purified by ethanol precipitation by adding 80 μL Milli-Q water, 10 μL sodium acetate (3 M), 1 μL GlycoBlue (Thermo Fisher Scientific #AM9515), and 250 μL ethanol. The purified DNA was then resuspended in 100 μL TE buffer and sheared using a Q800 sonicator (Qsonica) using the following program: 20% amplitude for 3 min, for 2 s on/5 s off. The fragmented DNA was subjected to size-selective purification to remove fragments <200 bp with 1x volume of AMPure XP DNA purification beads (Beckman Coulter) and eluted using 21 μL TE buffer. Eluted DNA was denatured by heating at 98 °C for 3 min, then immediately placed on ice. Next, 10 μL Dynabeads MyOne Streptavidin C1 (Thermo Fisher Scientific #65001) were washed twice with 1x BW buffer (5 mM Tris-HCl pH 8.0, 0.5 mM EDTA, 1 M NaCl, 0.1% Tween20), then resuspended with 20 μL 2 x BW buffer and added to the denatured DNA solution. The beads were rotated 1 h at 24 °C on a tube revolver with oscillation mode. Next, the beads were washed twice with LBW buffer (5 mM Tris-HCl pH 8.0, 0.5 mM EDTA, 150 mM NaCl, 0.1% Tween20) following with 3ʹ dephosphorylation by adding a solution containing 44.5 μL Milli-Q water, 5 μL 10 x CutSmart buffer (NEB #B7204), 0.5 μL dithiothreitol (0.5 M), and 1 μL T4 polynucleotide kinase (NEB #M0201). The mixture was incubated 30 min at 37 °C on a thermoshaker at 600 rpm and the tube was resuspended every 10 min in case of aggregation. After incubation, the beads were washed twice with LBW buffer and once with Milli-Q water. To elute DNA from beads, Milli-Q water (19 μL) was added to beads and the tube was incubated for 1 min at 98 °C. The supernatant was then transferred to a new 1.5 mL tube. The elution step was repeated once, and the supernatants were combined. The combined supernatant was then subjected to L-P7-5 adapter ligation by adding 5 μL 10 x T4 DNA ligase buffer, 5 μL 50% PEG-4000 solution, 0.5 μL 1% Tween 20, 2.5 μL L-P7-5 adapter (40 μM), and 0.5 μL T4 DNA Ligase (NEB, #M0202M). The mixture was incubated overnight (16 h) at 4 °C on a tube revolver with oscillation mode. The ligation mixture was then purified with 85 μL of ProNex Size-Selective Purification beads (Promega) and eluted with 30 μL 0.1 x TE buffer. To bypass the oxime linkage, the purified DNA was amplified for 5 cycles using Q5 High Fidelity DNA Polymerase with 0.5 μM of Ex-Bar-mix and L-P7-3end primers in 50 μL under the following condition: 60 s at 98 °C, then 10 s at 98 °C, 20 s at 59 °C, and 60 s at 72 °C for 5 cycles. The amplified product was purified using 85 μL ProNex beads and eluted by 30 μL 0.1 x TE buffer. The purified DNA product then can be used for qPCR quantification and undergo PCR with desired indexing primers for sequencing. The indexed libraries were sequenced as 1×100 on an Illumina NovaSeq6000 sequencer (Illumina).

### 5ʹ-AT sequencing data processing and analysis

Data quality was checked using FastQC and MultiQC,^43^ common adaptor contaminations were filtered and the 5 bases from read starts are trimmed using BBMap.^44^ The single-end reads were mapped to the reference human genome hg38 using Burrows-Wheeler Aligner.^45^ Unmapped and duplicated reads were removed using Samtools.^46^ The genomic coordinate of damage sites, which is the 1^st^ base of reads, were extracted and counted in desired bins using custom scripts and bedtools.^47^ The damage sites mapped within the ENCODE Blacklist^48^ were removed, then further analysis was either performed using DeepTools^49^, or using Bioconductor 3.1.2 under R 4.1.0 with custom scripts. Scaling factors were calculated using DESeq2^50^ and were applied to samples before analysis. DNase-seq,^51^ ATAC-seq, and ChIP-seq of histone mark^52,53^ data was obtained from previously published studies.

## Data Availability

The sequencing data underlying this article are available in the Sequence Read Archive (SRA) at https://www.ncbi.nlm.nih.gov/sra, and can be accessed with SRAXXXXXX.

## Reference

1 Cooke, M. S., Evans, M. D., Dizdaroglu, M. & Lunec, J. Oxidative DNA damage: mechanisms, mutation, and disease. Faseb J 17, 1195–1214 (2003). 10.1096/fj.02-0752rev

2 Markkanen, E. Not breathing is not an option: How to deal with oxidative DNA damage. DNA Repair (Amst) 59, 82–105 (2017). 10.1016/j.dnarep.2017.09.007

3 Caldecott, K. W. Single-strand break repair and genetic disease. Nat Rev Genet 9, 619–631 (2008). 10.1038/nrg2380

4 Dedon, P. C. The chemical toxicology of 2-deoxyribose oxidation in DNA. Chem Res Toxicol 21, 206–219 (2008). 10.1021/tx700283c

5 Balasubramanian, B., Pogozelski, W. K. & Tullius, T. D. DNA strand breaking by the hydroxyl radical is governed by the accessible surface areas of the hydrogen atoms of the DNA backbone. P Natl Acad Sci USA 95, 9738–9743 (1998). 10.1073/pnas.95.17.9738

6 Chan, W. et al. Quantification of the 2-deoxyribonolactone and nucleoside 5’-aldehyde products of 2-deoxyribose oxidation in DNA and cells by isotope-dilution gas chromatography mass spectrometry: differential effects of gamma-radiation and Fe2+-EDTA. J Am Chem Soc 132, 6145–6153 (2010). 10.1021/ja910928n

7 Boussicault, F., Kaloudis, P., Caminal, C., Mulazzani, Q. G. & Chatgilialoglu, C. The fate of C5’ radicals of purine nucleosides under oxidative conditions. J Am Chem Soc 130, 8377–8385 (2008). 10.1021/ja800763j

8 Rana, A., Yang, K. & Greenberg, M. M. Reactivity of the Major Product of C5’-Oxidative DNA Damage in Nucleosome Core Particles. Chembiochem 20, 672–676 (2019). 10.1002/cbic.201800663

9 Asaeda, A., Ide, H., Terato, H., Takamori, Y. & Kubo, K. Highly sensitive assay of DNA abasic sites in mammalian cells-optimization of the aldehyde reactive probe method. Analytica Chimica Acta 365, 35–41 (1998). 10.1016/S0003-2670(97)00648-X

10 Liu, Z. J., Martínez Cuesta, S., van Delft, P. & Balasubramanian, S. Sequencing abasic sites in DNA at single-nucleotide resolution. Nature Chemistry 11, 629–637 (2019). 10.1038/s41557-019-0279-9

11 Wu, W. et al. Neuronal enhancers are hotspots for DNA single-strand break repair. Nature 593, 440–444 (2021).

12 Cao, H. et al. Novel approach reveals genomic landscapes of single-strand DNA breaks with nucleotide resolution in human cells. Nature Communications 10, 5799 (2019). 10.1038/s41467-019-13602-7

13 Sriramachandran, A. M. et al. Genome-wide Nucleotide-Resolution Mapping of DNA Replication Patterns, Single-Strand Breaks, and Lesions by GLOE-Seq. Mol Cell 78, 975–985 e977 (2020). 10.1016/j.molcel.2020.03.027

14 Xu, S. et al. SSBlazer: a genome-wide nucleotide-resolution model for predicting single-strand break sites. Genome Biology 25, 46 (2024). 10.1186/s13059-024-03179-w

15 Kodama, T. & Greenberg, M. M. Preparation and analysis of oligonucleotides containing lesions resulting from C5’-oxidation. J Org Chem 70, 9916–9924 (2005). 10.1021/jo051666k

16 Holmberg, A. et al. The biotin-streptavidin interaction can be reversibly broken using water at elevated temperatures. Electrophoresis 26, 501–510 (2005). 10.1002/elps.200410070

17 Thrall, B. D., Mann, D. B., Smerdon, M. J. & Springer, D. L. Nucleosome structure modulates benzo[a]pyrenediol epoxide adduct formation. Biochemistry 33, 2210–2216 (1994). 10.1021/bi00174a030

18 Mann, D. B., Springer, D. L. & Smerdon, M. J. DNA damage can alter the stability of nucleosomes: effects are dependent on damage type. P Natl Acad Sci USA 94, 2215–2220 (1997). 10.1073/pnas.94.6.2215

19 Lambert, M. DNA repair mechanisms and their biological implications in mammalian cells. Vol. 182 (Springer Science & Business Media, 2013).

20 Poetsch, A. R., Boulton, S. J. & Luscombe, N. M. Genomic landscape of oxidative DNA damage and repair reveals regioselective protection from mutagenesis. Genome Biol 19, 215 (2018). 10.1186/s13059-018-1582-2

21 Mercer, T. R. et al. The human mitochondrial transcriptome. Cell 146, 645–658 (2011). 10.1016/j.cell.2011.06.051

22 Becker, J. S. et al. Genomic and Proteomic Resolution of Heterochromatin and Its Restriction of Alternate Fate Genes. Mol Cell 68, 1023-1037.e1015 (2017). 10.1016/j.molcel.2017.11.030

23 Maurano, M. T. et al. Large-scale identification of sequence variants influencing human transcription factor occupancy in vivo. Nat Genet 47, 1393–1401 (2015). 10.1038/ng.3432

24 Donaghey, J. et al. Genetic determinants and epigenetic effects of pioneer-factor occupancy. Nat Genet 50, 250–258 (2018). 10.1038/s41588-017-0034-3

25 Koch, C. M. et al. The landscape of histone modifications across 1% of the human genome in five human cell lines. Genome Res 17, 691–707 (2007). 10.1101/gr.5704207

26 Clarkson, C. T. et al. CTCF-dependent chromatin boundaries formed by asymmetric nucleosome arrays with decreased linker length. Nucleic Acids Res 47, 11181–11196 (2019). 10.1093/nar/gkz908

27 Cheng, D. et al. The genome-wide transcriptional regulatory landscape of ecdysone in the silkworm. Epigenetics Chromatin 11, 48 (2018). 10.1186/s13072-018-0216-y

28 Chen, H. et al. An integrative analysis of TFBS-clustered regions reveals new transcriptional regulation models on the accessible chromatin landscape. Sci Rep 5, 8465 (2015). 10.1038/srep08465

29 Soufi, A., Donahue, G. & Zaret, K. S. Facilitators and impediments of the pluripotency reprogramming factors’ initial engagement with the genome. Cell 151, 994–1004 (2012). 10.1016/j.cell.2012.09.045

30 Guo, Q. et al. How far can hydroxyl radicals travel? An electrochemical study based on a DNA mediated electron transfer process. Chemical Communications 47, 11906–11908 (2011). 10.1039/C1CC14699H

31 Henle, E. S. et al. Sequence-specific DNA Cleavage by Fe^2+^-mediated Fenton Reactions Has Possible Biological Implications *. Journal of Biological Chemistry 274, 962–971 (1999). 10.1074/jbc.274.2.962

32 Wang, W., Lee, G. J., Jang, K. J., Cho, T. S. & Kim, S. K. Real-time detection of Fe.EDTA/H2O2-induced DNA cleavage by linear dichroism. Nucleic Acids Res 36, e85 (2008). 10.1093/nar/gkn370

33 Cadet, J., Douki, T., Gasparutto, D. & Ravanat, J. L. Oxidative damage to DNA: formation, measurement and biochemical features. Mutat Res 531, 5–23 (2003). 10.1016/j.mrfmmm.2003.09.001

34 Wu, J., Sturla, S. J., Burrows, C. J. & Fleming, A. M. Impact of DNA Oxidation on Toxicology: From Quantification to Genomics. Chemical Research in Toxicology 32, 345–347 (2019). 10.1021/acs.chemrestox.9b00046

35 Nabbi, A. & Riabowol, K. Isolation of Nuclei. Cold Spring Harb Protoc 2015, 731–734 (2015). 10.1101/pdb.top074583

36 Ernst, J. et al. Mapping and analysis of chromatin state dynamics in nine human cell types. Nature 473, 43–49 (2011). 10.1038/nature09906

37 Ma, Y., Kanakousaki, K. & Buttitta, L. How the cell cycle impacts chromatin architecture and influences cell fate. Front Genet 6, 19 (2015). 10.3389/fgene.2015.00019

38 Collins, A. R., Ma, A. G. & Duthie, S. J. The kinetics of repair of oxidative DNA damage (strand breaks and oxidised pyrimidines) in human cells. Mutat Res 336, 69–77 (1995). 10.1016/0921-8777(94)00043-6

39 Frankenberg-Schwager, M. Review of repair kinetics for DNA damage induced in eukaryotic cells in vitro by ionizing radiation. Radiother Oncol 14, 307–320 (1989). 10.1016/0167-8140(89)90143-6

40 Hu, J., Adar, S., Selby, C. P., Lieb, J. D. & Sancar, A. Genome-wide analysis of human global and transcription-coupled excision repair of UV damage at single-nucleotide resolution. Genes Dev 29, 948–960 (2015). 10.1101/gad.261271.115

41 Motorin, Y. & Helm, M. General Principles and Limitations for Detection of RNA Modifications by Sequencing. Acc Chem Res 57, 275–288 (2024). 10.1021/acs.accounts.3c00529

42 Nabbi, A. & Riabowol, K. Rapid Isolation of Nuclei from Cells In Vitro. Cold Spring Harb Protoc 2015, 769–772 (2015). 10.1101/pdb.prot083733

43 Ewels, P., Magnusson, M., Lundin, S. & Kaller, M. MultiQC: summarize analysis results for multiple tools and samples in a single report. Bioinformatics 32, 3047–3048 (2016). 10.1093/bioinformatics/btw354

44 Bushnell, B., Rood, J. & Singer, E. BBMerge - Accurate paired shotgun read merging via overlap. PLoS One 12, e0185056 (2017). 10.1371/journal.pone.0185056

45 Li, H. & Durbin, R. Fast and accurate short read alignment with Burrows-Wheeler transform. Bioinformatics 25, 1754–1760 (2009). 10.1093/bioinformatics/btp324

46 Danecek, P. et al. Twelve years of SAMtools and BCFtools. Gigascience 10 (2021). 10.1093/gigascience/giab008

47 Quinlan, A. R. & Hall, I. M. BEDTools: a flexible suite of utilities for comparing genomic features. Bioinformatics 26, 841–842 (2010). 10.1093/bioinformatics/btq033

48 Amemiya, H. M., Kundaje, A. & Boyle, A. P. The ENCODE Blacklist: Identification of Problematic Regions of the Genome. Sci Rep 9, 9354 (2019). 10.1038/s41598-019-45839-z

49 Ramirez, F. et al. deepTools2: a next generation web server for deep-sequencing data analysis. Nucleic Acids Res 44, W160–165 (2016). 10.1093/nar/gkw257

50 Love, M. I., Huber, W. & Anders, S. Moderated estimation of fold change and dispersion for RNA-seq data with DESeq2. Genome Biol 15, 550 (2014). 10.1186/s13059-014-0550-8

51 Thurman, R. E. et al. The accessible chromatin landscape of the human genome. Nature 489, 75–82 (2012). 10.1038/nature11232

52 Consortium, E. P. An integrated encyclopedia of DNA elements in the human genome. Nature 489, 57–74 (2012). 10.1038/nature11247

53 Davis, C. A. et al. The Encyclopedia of DNA elements (ENCODE): data portal update. Nucleic Acids Res 46, D794–D801 (2018). 10.1093/nar/gkx1081

